# The mitochondrial genomes of the acoelomorph worms *Paratomella rubra* and *Isodiametra pulchra*

**DOI:** 10.1101/103556

**Authors:** Helen E Robertson, François Lapraz, Bernhard Egger, Maximilian J Telford, Philipp H. Schiffer

## Abstract

Acoels are small, ubiquitous, but understudied, marine worms with a very simple body plan. Their internal phylogeny is still in parts unresolved, and the position of their proposed phylum Xenacoelomorpha (Xenoturbella+Acoela) is still debated.

Here we describe mitochondrial genome sequences from two acoel species: *Paratomella rubra* and *Isodiametra pulchra*. The 14,954 nucleotide-long *P. rubra* sequence is typical for metazoans in size and gene content. The larger *I. pulchra* mitochondrial genome contains both ribosomal genes, 21 tRNAs, but only 11 protein-coding genes. We find evidence suggesting a duplicated sequence in the *I. pulchra* mitochondrial genome.

Mitochondrial sequences for both *P. rubra* and *I. pulchra* have a unique genome organisation in comparison to other published metazoan mitochondrial genomes. We found a large degree of protein-coding gene and tRNA overlap in *P. rubra*, with little non-coding sequence making the genome compact. Conversely, the *I. pulchra* mitochondrial genome has many long non-coding sequences between genes, likely driving the genome size expansion. Phylogenetic trees inferred from concatenated alignments of mitochondrial genes grouped the fast-evolving Acoela and Tunicata, almost certainly due to the systematic error of long branch attraction: a reconstruction artefact that is probably compounded by the fast substitution rate of mitochondrial genes in this taxon.

## Introduction

Acoel flatworms are small, soft-bodied, unsegmented, marine animals lacking a gut epithelium, coelomic cavity, and anus. Instead, they typically possess a ventral mouth opening, and a simple syncytial digestive system^1^. Due primarily to the common attributes of acoelomate body and the absence of a through gut, Acoela were traditionally grouped as an order within the Platyhelminthes. The first molecular systematic studies on these animals using small subunit (SSU) ribosomal RNA gene sequences revealed that the Acoelomorpha are in fact a distinct lineage, quite separate from the main clade of the Platyhelminthes (Rhabditophora and Catenulida) ^2–4^. Instead, these initial molecular studies supported a position of the Acoelomorpha diverging prior to the protostome/deuterostome common ancestor. More recently, the Acoelomorpha have been linked to the similarly simple marine worm *Xenoturbella* in the new phylum Xenacoelomorpha, making sense of their shared simple body plan and other shared morphological characters, such as unusual ciliary ultrastructure^5^ and their simple basiepidermal nervous system^6^. Despite considerable efforts, the position of Xenacoelomorpha within the Metazoa remains unresolved, with alternative lines of evidence placing them either as the sister group to the remaining Bilateria (protostomes and deuterostomes)^7,8^, or as a phylum within the deuterostomes^9^. A better understanding of acoel phylogeny and evolution is therefore integral to answering central questions concerning the evolution of Bilateria and its subtaxa. To this end more genomic data are needed.

Metazoan mitochondrial DNA (mtDNA) is a closed-circular molecule typically comprising 37 genes which are, for the most part, invariant across the Metazoa^10^. These include the two rRNAs of the mitochondrial ribosome, 22 tRNAs necessary for translation, and 13 protein-coding genes for the enzymes of oxidative phosphorylation. *atp8* is the only gene known to have been commonly lost from this complement, and this has been observed in a number of independent metazoan lineages, including the acoel *Symsagittifera roscoffensis*^11^. In addition to primary sequence data, mtDNA has a number of other features which can be used for phylogenetic inference, including variations in mitochondrial genetic code^12^; a higher rate of sequence evolution than nuclear DNA^13^; changes in gene order and changes in the secondary structure of rRNAs and tRNAs^14^.

Mitochondrial gene sequences have been used extensively for phylogenetic inference. In a recent paper, Rouse et al. used mitochondrial protein-coding sequence data from four newly discovered species of *Xenoturbella* (*X. hollandorum*, *X. churro*, *X. monstrosa* and *X. profunda*) to infer the internal phylogeny of the Xenoturbellida^15^. Wider phylogenetic inference including mitochondrial proteins from these species placed Xenacoelomorpha with the deuterostomes^15^, corroborating previous mitochondrial phylogenetic analysis of this phylum^9,16,17^.

Mitochondrial gene content is largely invariable across the Metazoa, with the order in which genes are arranged being fairly stable and conserved for up to hundreds of millions of years in some metazoan lineages. Rearrangement events, thought to occur via a model of ‘duplication and deletion’^14,18^, whereby a portion of the mitochondrial genome is duplicated, and the original copy of the duplicated gene subsequently deleted, are rare. The infrequency of such rearrangements, and the huge number of possible rearrangement scenarios, means that convergence on the same gene order in unrelated lineages is unlikely. Gene order is thus likely to retain evolutionary signals, with a common gene order being indicative of common ancestry and informative for the study of metazoan divergence^19^. Rearrangement of genes within the mitochondrial genome of different species can be a particularly powerful tool in the analysis of phylogenetic relationships^14^ and may also indicate accelerated evolution in a taxon.

In this study, we describe the mitochondrial genomes from two species of acoel: *Paratomella rubra* and *Isodiametra pulchra*. Adult specimens of both animals are approximately 1mm in length, and, as is typical for small acoel species, they occupy the littoral and sub-littoral zones of marine ecosystems: *P. rubra* has been described across Europe and North America^20,21^, and *I. pulchra* lives abundantly in the mud flats of Maine^22^. Both species move freely within the sediment by gliding on a multiciliated epidermis. First described by Rieger and Ott^21^, *P. rubra* is an elongate and flattened worm belonging to the family Paratomellidae, which is sister group to all other described acoel species^23,24^. A 9.7kb fragment of mitochondrial genome has previously been described from specimens of *P. rubra* collected on the Mediterranean coast of Spain^25^. *I pulchra* belongs to the family Isodiametridae; it can be maintained long-term in culture and has been used experimentally for *in situ* hybridisation, RNAi, and other molecular protocols^22,26,27^. It's use as a ‘model acoel’ therefore makes this species particularly valuable for investigation.

## Results

### Genomic Composition

We assembled 14,954 base pairs of the *P. rubra* mitochondrial genome, starting from three genome assembly fragments and using Sanger sequencing of PCR fragments (Fig. 1a). We were unable to close the circular mitochondrial genome of *P. rubra*, but our 14.9kb sequence contains all 13 protein-coding genes, both ribosomal genes and 22 putative tRNAs. Compared to the fragment of the genome previously published we have found four additional protein-coding genes and 12 additional tRNAs ^25^. All genes are found exclusively on one strand of the sequence. Allowing for overlap, protein-coding genes account for 74.79% of the genomic sequence; ribosomal genes 13.95%; tRNA genes 9.10% and non-coding DNA 2.04%. A 156 nucleotide-long stretch of non-coding sequence is found between *cytochrome c oxidase subunit 2* (*cox2*) and *NADH dehydrogenase subunit 1* (*nad1*).

**Figure 1:**
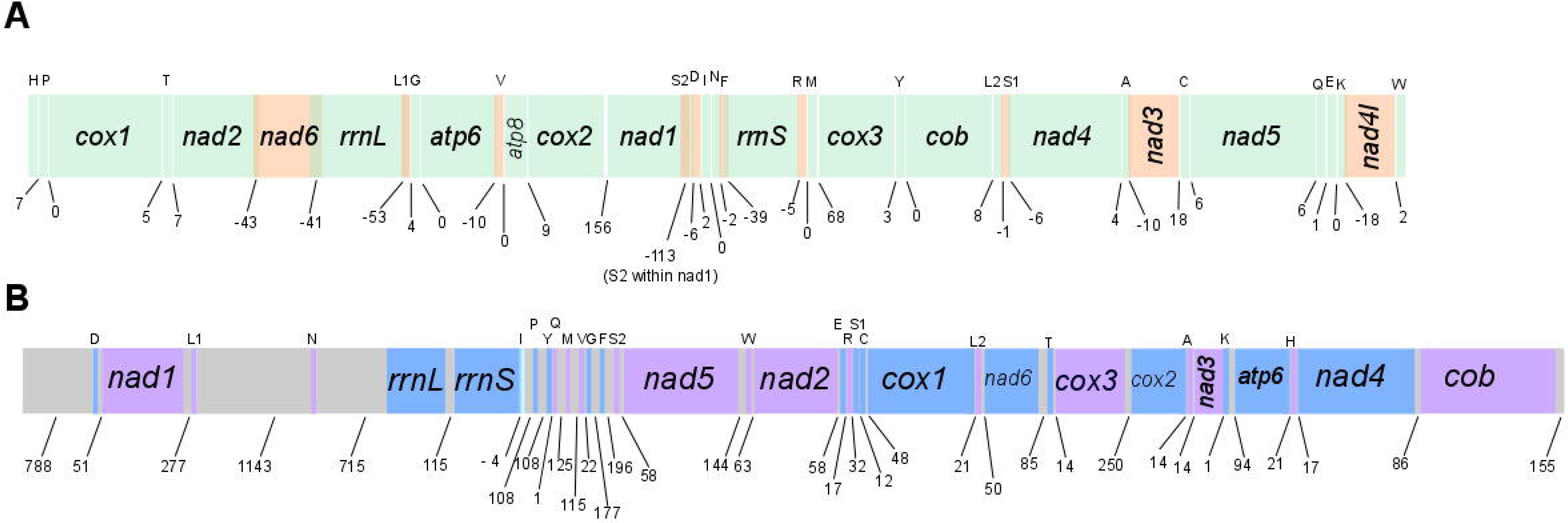
Overview of the mitochondrial genome sequences we resolve for *Paratomella rubra* and *Isodiametra pulchra* (Xenacoelomorpha: Acoela). Genes not drawn to scale. Numbers beneath the sequences show intergenic spaces (positive values) or intergenic overlap (negative values). Protein-coding genes are denoted by three letter abbreviations; ribosomal genes by four letter abbreviations. tRNAs are shown by single uppercase letters. **A**. *P. rubra* 14,957 base-pair ling sequence. All genes found on the positive (forward) strand. Where genes, rRNAs or tRNAs are coloured orange, this is solely to demonstrate overlap with adjacent genes, rRNAs or tRNAs. **B**. *I. pulchra* 18, 725 base-pair long sequence. Genes found on the positive (forward) strand are coloured blue; genes on the negative (reverse) strand are coloured purple. Non-coding sequence shown in grey.

In *P. rubra*, *trn*S2 is predicted entirely within the sequence coding for *nad1*, and also has clear deviation from the traditional 'cloverleaf' secondary structure of tRNA. In addition, three of the predicted tRNAs have minor overlap with protein-coding genes: *trn*A with *nad3* (20 nucleotides); *trnK* with *nad4l* (18 nucleotides) and *trnS1* with *nad4* (six nucleotides); and all but five nucleotides of *trnL1* are predicted within the same sequence as *rrnL* (Fig. 1, Table 1). With the exception of *trnT*, all predicted tRNAs have an amino-acyl acceptor stem composed of seven base pairs, and all predicted tRNAs apart from *trnT* and *trnS2* have a five base pair anticodon stem (Fig. 3). 11 *tRNAs* have one or two G-T mismatches in their acceptor or anticodon stems (*A,C,G,I,K,L1,L2,P,Q,R,T*). All tRNAs have a DHU arm of three or four nucleotides. The structure of the TψC arm shows greater variability, with a number of tRNAs having either a truncated stem, or the arm entirely lacking (Fig. 3).

**Table 1:** Organisation of the *Paratomella rubra* 14.9kb mitochondrial genome sequence. All genes found on the 'positive' strand

| Gene | Start | End | Length (bp) | Length (amino acid) | Start Codon | Stop Codon | Intergenic nucleotides |
| --- | --- | --- | --- | --- | --- | --- | --- |
| *trnH*(gtg) | 368 | 426 | 59 |  |  |  | 7 |
| *trnP*(tgg) | 434 | 495 | 62 |  |  |  | 0 |
| *Cox1* | 496 | 2058 | 1563 | 521 | ATA | TAA | 5 |
| *trnT*(tgt) | 2064 | 2123 | 54 |  |  |  | 7 |
| *Nad2* | 2131 | 3144 | 1014 | 338 | ATT | TAG | -43 |
| *Nad6* | 3102 | 3563 | 462 | 154 | ATA | TAA | -101 |
| *rrnL* | 3463 | 4819 | 1357 |  |  |  | -53 |
| *trnL1*(tag) | 4767 | 4824 | 58 |  |  |  | 4 |
| *trnG*(tcc) | 4829 | 4887 | 59 |  |  |  | 0 |
| *Atp6* | 4888 | 5496 | 609 | 203 | ATA | TAG | -10 |
| *trnV*(tac) | 5487 | 5553 | 67 |  |  |  | 0 |
| *Atp8* | 5554 | 5730 | 177 | 59 | ATT | TA- | 9 |
| *Cox2* | 5740 | 6402 | 663 | 221 | ATT | TAA | **156** |
| *Nad1* | 6559 | 7611 | 1053 | 351 | ATT | TAA | -113 |
| *trnS2*(tga) | 7499 | 7568 | 70 |  |  |  | -6 |
| *trnD*(gtc) | 7601 | 7662 | 62 |  |  |  | 2 |
| *trnI*(gat) | 7665 | 7728 | 64 |  |  |  | 0 |
| *trnN*(gtt) | 7729 | 7798 | 70 |  |  |  | -2 |
| *trnF*(gaa) | 7797 | 7856 | 60 |  |  |  | -39 |
| *rrnS* | 7818 | 8547 | 730 |  |  |  | -5 |
| *trnR*(tcg) | 8542 | 8608 | 67 |  |  |  | 0 |
| *trnM*(cat) | 8609 | 8669 | 61 |  |  |  | 68 |
| *Cox3* | 8738 | 9523 | 786 | 262 | ATT | TAA | 3 |
| *trnY*(gta) | 9527 | 9585 | 59 |  |  |  | 0 |
| *Cob* | 9586 | 10668 | 1083 | 361 | ATA | TAA | 8 |
| *trnL2*(taa) | 10677 | 10737 | 61 |  |  |  | -1 |
| *trnS1*(gct) | 10736 | 10799 | 64 |  |  |  | -6 |
| *Nad4* | 10794 | 12119 | 1326 | 442 | ATC | TAA | 4 |
| *trnA*(tgc) | 12124 | 12181 | 58 |  |  |  | -10 |
| *Nad3* | 12162 | 12551 | 390 | 130 | ATT | TAG | 18 |
| *trnC*(gca) | 12570 | 12629 | 60 |  |  |  | 6 |
| *Nad5* | 12636 | 14387 | 1752 | 584 | ATA | TAG | 6 |
| *trnQ*(ttg) | 14394 | 14449 | 56 |  |  |  | 1 |
| *trnE*(ttc) | 14451 | 14510 | 60 |  |  |  | 0 |
| *trnK*(ttt) | 14511 | 14576 | 66 |  |  |  | -18 |
| *Nad4l* | 14559 | 14867 | 309 | 103 | ATA | TAA | 2 |
| *trnW*(tca) | 14870 | 14933 | 64 |  |  |  |  |

**Figure 3:**
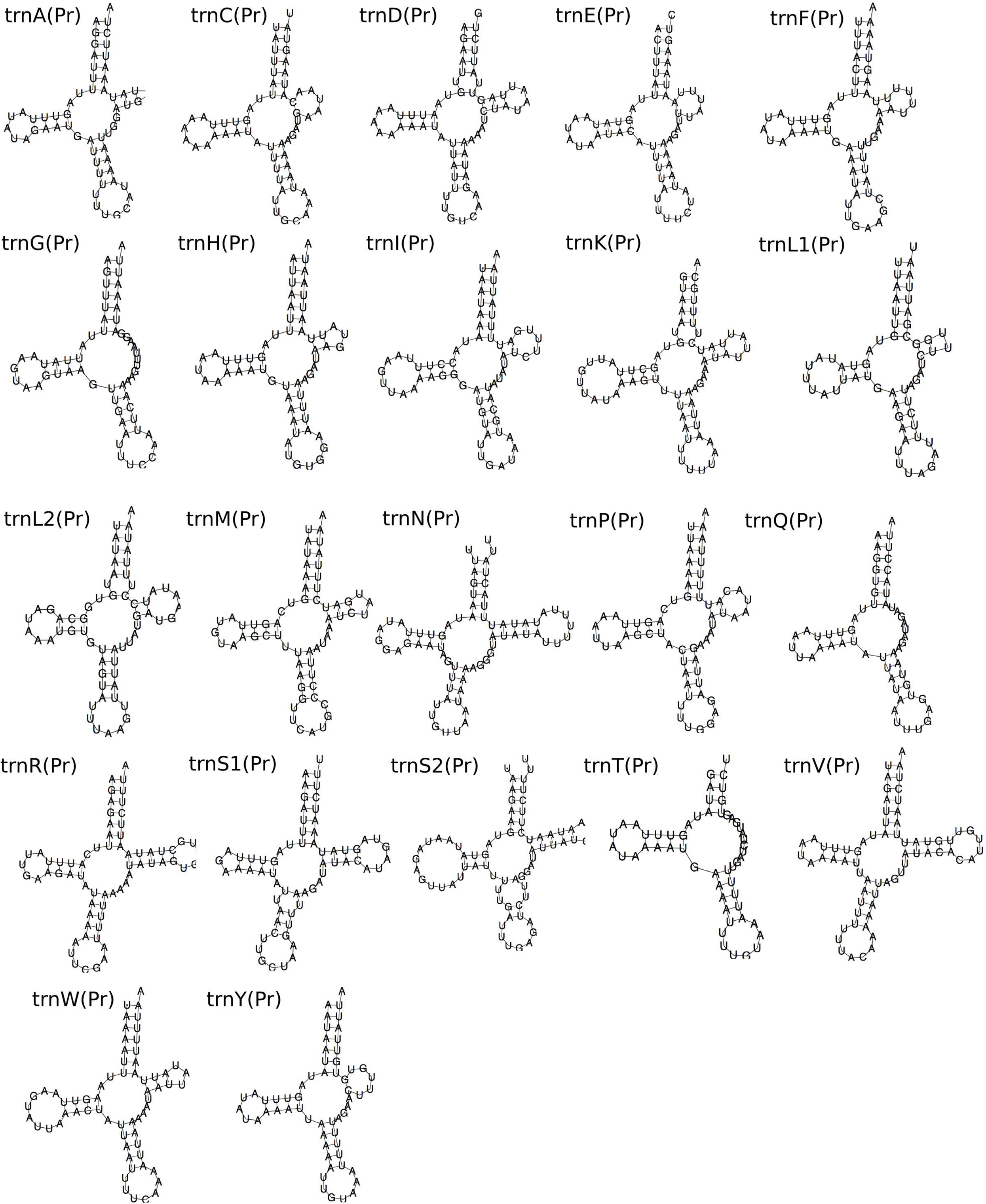
Predicted secondary structures of tRNAs from the mitochondrial genome sequence of *Paratomella rubra* as predicted by MiTFi in Mitos.

For *I. pulchra*, we initially recovered a 13kb fragment, a 3.5kb fragment and a 19kb fragment of mitochondrial sequence from our transcriptomic data. The entire 13kb fragment and 2.4kb of the 3.5kb fragment were found to be perfectly matching subsets of the longer 19kb sequence (Fig. 2). We designed several sets of PCR primers to verify the sequence between the 3’ end of the 13kb and 5’ end of the 3.5kb fragments found on the long 19kb fragment (Fig. 1b), however, no PCR amplification completely bridged the sequence between the 13kb to 3.5kb fragment. We found that the last (3’) 300bp of the 13kb fragment was duplicated in the opposite orientation within the end (3’) region of the 3.5kb fragment. Although the long 19kb fragment indicated that the repeated region between the 13kb and 3.5kb fragments was present, we were not able to connect sequence on both sides of the repeated region by PCR. The placement of Sanger sequencing fragments containing the repeat remained ambiguous. We are thus not confident in the assembly of the 19kb transcriptomic sequence in this section, and therefore treat it as unresolved. Instead we focused on verifying the sequence of the two smaller fragments and on amplifying and sequencing the region lying between them. We reconfirmed the majority of the 13kb fragment using PCR amplification and Sanger sequencing. We were also able to amplify and sequence fragments joining the 3’ end of the 3.5kb fragment with the 5’ end of a 1.3kb fragment containing the *rrnL* gene, which we had identified in transcriptomic data with tblastn. This included the occurrence of the duplicated region at the 3’ end of the 3.5kb fragment (Fig. 2).

**Figure 2:**
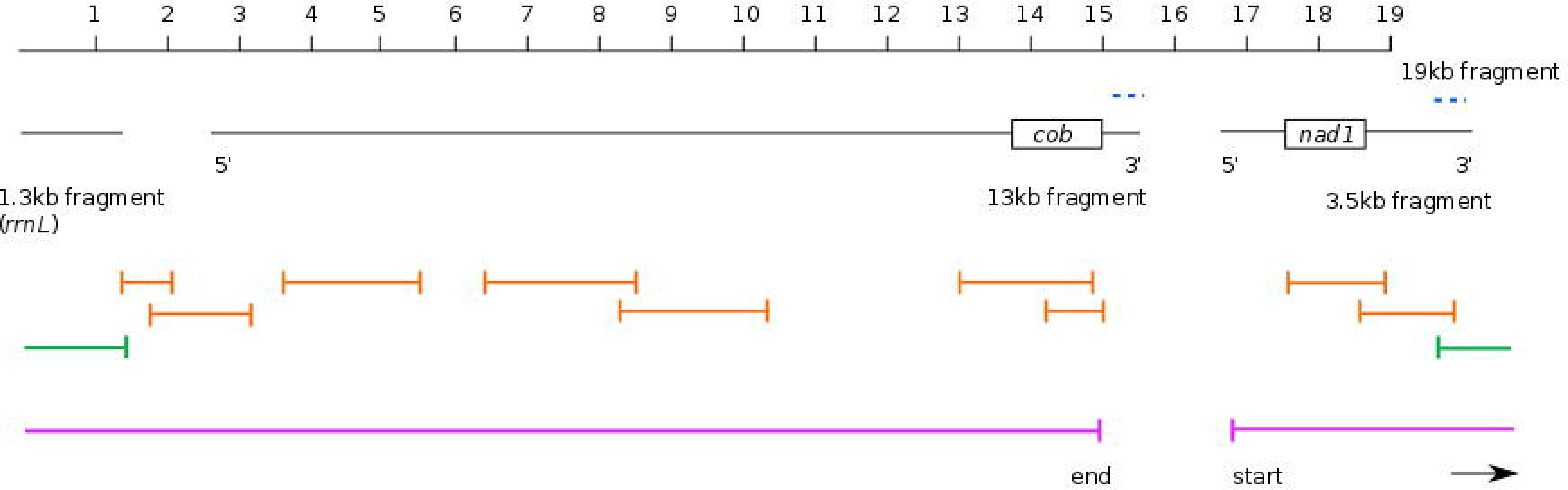
Overview of the initial transcriptome assembly fragments and PCR strategy for scaffolding the *Isodiametra pulchra* mitochondrial genome. 1.3kb, 13kb and 3.5kb fragments aligned to continuous 19kb fragment, with the location of the duplicated sequence in the 13kb and 3.5kb fragments shown by blue dashed lines. The ‘start’ and ‘end’ regions of the 13kb and 3.5kb fragments are annotated by 5’ (start) and 3’ (end). The approximate location of *cob* and *nad1* protein-coding sequence are shown for reference. Reliable PCR-amplicons are shown in orange; the green PCR fragment indicates successful joining of the 3’ end of the 3.5kb fragment to the *rrnL* fragment, including the duplicated section. The 18, 725 base-pair long sequence we resolve is indicated by the pink lines, from ‘start’ to ‘end’.

In summary, we find the *I. pulchra* mitochondrial genome to have a span of at least 18,725 base pairs (Fig. 1b) based on our PCR validation of the transcriptomic data. This covers the region from the start of the 5’ end of the 3.5kb sequence which is linked through PCR amplicons to the 5’ end of the 13kb sequence (including the 1.3kb *rrnL* contig), and up to the start of the duplicated sequence at the 3’ end of the 13kb sequence (Fig. 2). As we were not able to bridge the region between the 3’ end of the 13kb fragment and the 5’ end of the 3.5kb fragment with PCR, we could not confirm the validity of the duplicated sequence at this position nor fully close the circular mitochondrial genome. It is therefore likely that the entire mitochondrial genome is larger than 19kb, and may include the duplicated sequence. The sequence we are confident on presenting contains both ribosomal genes, all tRNAs and 11 protein-coding genes. These protein-coding genes and RNAs are encoded on both the plus and minus strands. No sequences resembling either *atp8* or *nad4l* could be found in our sequence.

In the 18.7kb sequence, protein-coding genes account for 56.66%; ribosomal genes contribute 8.15% and tRNA genes 7.77%. Compared to *the P.rubra* and *S. roscoffensis* mitochondrial genomes, intergenic space in the *I. pulchra* sequences is unusually high: non-coding DNA accounts for 22.72% of the sequences, including 13 intergenic regions of greater than 100 base pairs.

We identified all 22 expected tRNAs in the *I. pulchra* mitochondrial genome. Predicted sequences for *rrnS* and *trnI* overlap by four base pairs, but no other overlaps were found between any tRNAs or with any protein-coding genes (Fig. 1b, Table 2). All predicted tRNAs have an amino-acyl acceptor stem composed of seven base pairs and a five base pair anticodon stem, with the exception of *trnE, trnF* and *trnS2*, which have an anticodon stem composed of only four base pairs (Fig. 4). The structure of the DHU arms and TψC show greater variability, and are composed of either 3 or 4, or between 3 and 6, base pairs respectively, across the 22 tRNAs. Whilst the TψC arm is missing entirely in *trnQ*, and very truncated in *trnE*, *trnF*, *trnG* and *trnP*, more of the predicted tRNAs fit the stereotypical 'cloverleaf' secondary structure than has been found for other acoel species, including *S. roscoffensis* and *P. rubra* (Fig. 4).

**Table 2:** Organisation of the *Isodiametra pulchra* 18.7kb mitochondrial genome sequence.

| Gene | Strand | Start | End | Length (bp) | Length (amino acid) | Start Codon | Stop Codon | Intergenic nucleotides |
| --- | --- | --- | --- | --- | --- | --- | --- | --- |
| *trnD* (gtc) | + | 789 | 848 | 68 |  |  |  | 51 |
| *Nad1* | - | 900 | 1784 | 885 | 295 | ATG | TAA | 277 |
| *trnL1* (tag) | - | 2062 | 2130 | 69 |  |  |  | 1143 |
| *trnN* (gtt) | - | 3274 | 3339 | 64 |  |  |  | 717 |
| *rrnL* | + | 4057 | 4657 | 601 |  |  |  | 115 |
| *rrnS* | + | 4773 | 5698 | 926 |  |  |  | -4 |
| *trnI* (gat) | + | 5695 | 5768 | 74 |  |  |  | 108 |
| *trnP* (tgg) | + | 5877 | 5939 | 63 |  |  |  | 108 |
| *trnY* (gta) | + | 6048 | 6111 | 64 |  |  |  | 1 |
| *trnQ* (ttg) | - | 6113 | 6173 | 61 |  |  |  | 125 |
| *trnM* (cat) | - | 6299 | 6360 | 62 |  |  |  | 115 |
| *trnV* (tac) | - | 6476 | 6543 | 68 |  |  |  | 22 |
| *trnG* (tcc) | + | 6566 | 6627 | 62 |  |  |  | 177 |
| *trnF* (gaa) | + | 6805 | 6873 | 69 |  |  |  | 196 |
| *trnS2* (tga) | - | 7070 | 7139 | 70 |  |  |  | 58 |
| *Nad5* | - | 7198 | 8907 | 1710 | 570 | ATG | TAA | 144 |
| *trnW* | - | 9052 | 9118 | 67 |  |  |  | 63 |
| *Nad2* | - | 9181 | 10233 | 1053 | 351 | ATG | TAA | 58 |
| *trnE* (ttc) | + | 10292 | 10355 | 64 |  |  |  | 17 |
| *trnR* (tcg) | - | 10373 | 10439 | 67 |  |  |  | 32 |
| *trnS1* (tct) | + | 10472 | 10539 | 68 |  |  |  | 12 |
| *trnC* (gca) | + | 10551 | 10613 | 63 |  |  |  | 48 |
| *Cox1* | + | 10662 | 12197 | 1536 | 512 | ATA | TAG | 21 |
| *trnL2* (taa) | - | 12219 | 12286 | 68 |  |  |  | 50 |
| *Nad6* | + | 12336 | 12811 | 476 | 159 | ATG | T-- | 85 |
| *trnT* (tgt) | + | 12897 | 12963 | 67 |  |  |  | 14 |
| *Cox3* | - | 12978 | 13775 | 798 | 266 | ATG | TAA | 250 |
| *Cox2* | + | 14026 | 14640 | 615 | 205 | ATA | TAA | 14 |
| *trnA* (tgc) | - | 14655 | 14718 | 64 |  |  |  | 14 |
| *Nad3* | - | 14733 | 15110 | 378 | 126 | ATG | TAA | 1 |
| *trnK* (ttt) | + | 15112 | 15178 | 67 |  |  |  | 94 |
| *Atp6* | + | 15273 | 15955 | 681 | 229 | ATA | TA- | 21 |
| *trnH* (gtg) | - | 15977 | 16042 | 66 |  |  |  | 17 |
| *Nad4* | + | 16060 | 17403 | 1344 | 448 | ATG | TAA | 86 |
| *Cob* | - | 17490 | 18570 | 1134 | 378 | ATA | T-- |  |

**Figure 4:**
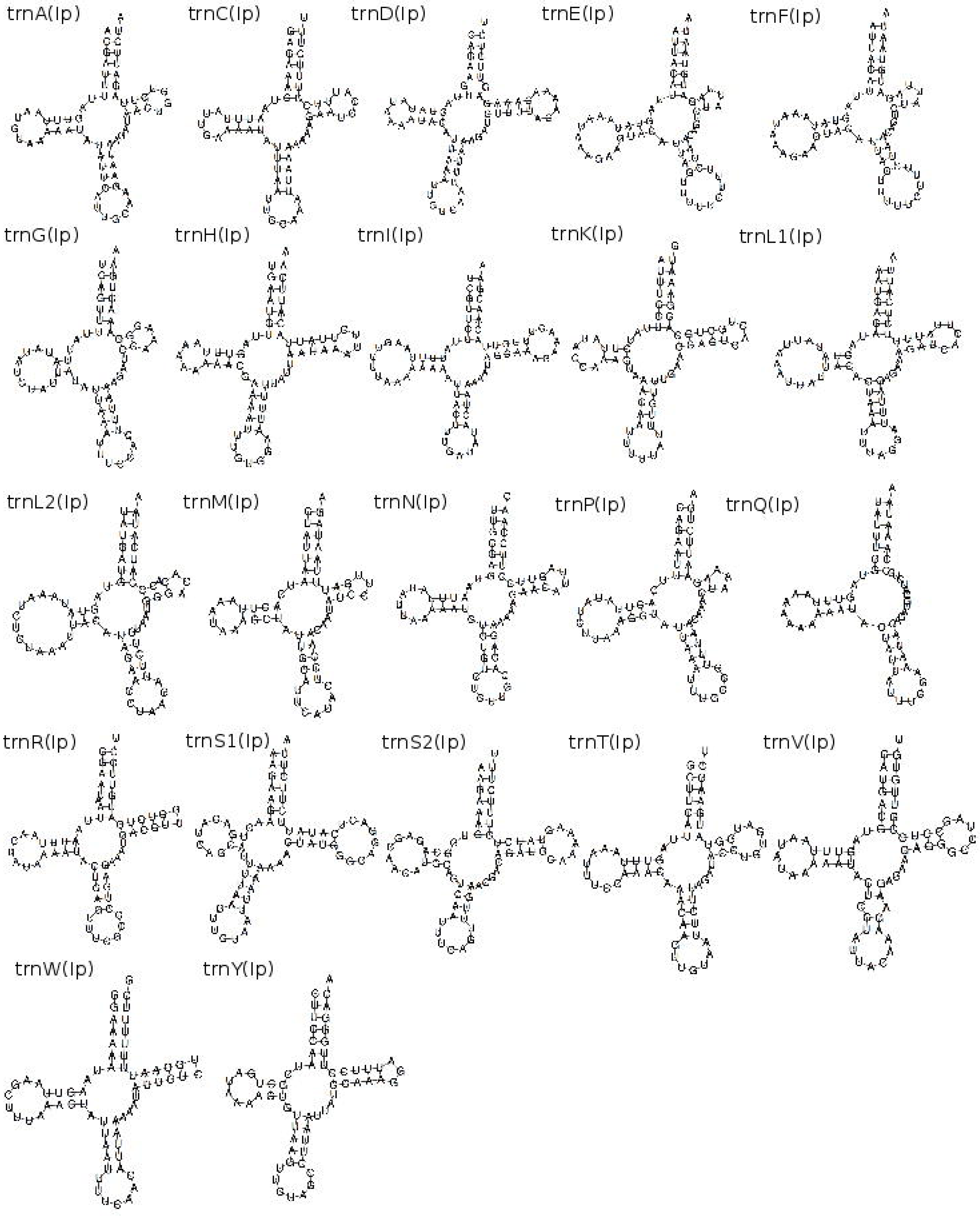
Predicted secondary structure of tRNAs from the mitochondrial genome sequences of and *Isodiametra pulchra* as predicted by MiTFi in Mitos. *trnN*, predicted within the duplicated sequence, shown in red.

The *P. rubra* genome is 78.15% A+T rich, which is higher than the A+T content calculated for the *I. pulchra* genomic fragments at 67.28%. Overall nucleotide usage on the plus strand of *P. rubra* is: A = 29.29%; T = 48.86%; C = 6.77% and G = 15.10%; GC-skew = 0.38 and absolute AT-skew = 0.25. Overall nucleotide usage for *I. pulchra* is: A = 34.04%; T = 33.24%; C = 16.45% and G = 16.27%; GC-skew = 0.006 and AT-skew = 0.012. GC-skew and AT-skew absolute values for *P. rubra* are much higher than that of *S. roscoffensis*, although the absolute values for *I. pulchra* are comparatively low (0.02 and 0.05 respectively)^11^. AT-skew value for the *P. rubra* sequence is just 0.01 different from that of the published partial *P. rubra* genome, and GC-skew is slightly higher (published *P. rubra* GC-skew = 0.32)^25^.

### Gene Order and Arrangement

All thirteen protein-coding genes in *P. rubra* have complete initiation codons: ATA (x5) and ATT (x8). Five of the protein-coding genes previously published differ in the nucleotide sequence of their start codons: *nad2*, *atp8*, *cox2*, and *cox3* all have ATA as an initiation codon in our analysis, compared to ATT found in previous analysis^25^. Twelve of the genes have full stop codons: TAA (x9) or TAG (x3). *atp8* was found to have a truncated stop codon (TA-), which is thought to be completed during post-transcriptional modification (Table 1). The eleven protein-coding genes found for *I. pulchra* also have full initiation codons: ATA (x4) and ATG (x7). Eight of the genes for this species have full stop codons: TAA (x7) and TAG (x1); *nad6*, *atp6* and *cob* are inferred to have truncated stop codons (Table 2). As in other invertebrate mitochondrial genomes, our data indicates a deviation from the 'standard' genetic code, with ATA encoding the start codon methionine, M, instead of isoleucine, I.

We found all genes in *P. rubra* on the ‘plus’ strand. In *I. pulchra*, genes are distributed over the plus and minus strands, with just two ‘blocks’ of genes with the same transcriptional polarity clustered together (*rrnL-rrnS-trnI-trnP-trnY*; *trnS2-nad5-trnW-nad2*). Whilst the *P. rubra* mitochondrial sequence has a large degree of overlap between adjacent genes, the opposite is true for *I. pulchra*: there are 4 base pairs of coding–sequence overlap across the two consensus sequences, between *rrnL* and *rrnS*. Unlike other metazoan mitochondrial genomes, where genes are adjacent or overlapping and one or two larger non-coding regions are commonly found, *I. pulchra* non-coding sequence is found consistently between protein-coding genes and between tRNAs, ranging in length from twelve to 277 base pairs. In addition, three long non-coding regions of 788, 1143 and 717 base pairs are found at the start of our sequence; between *trnL1* and *trnN*; and *trnN* and *rrnL*. The A+T content of these three sections are 68.78%, 65.79% and 76.15% respectively. The compositional difference between the 717 base pair non-coding region and the rest of the genome is statistically different (χ2 = 25.629, p<0.0001), with a higher A+T content indicating that it could function as a transcriptional control region. The A+T content of the 788 base pair non-coding region is not significantly higher than the rest of the sequence (χ2 = 0.850). The *P. rubra* sequence has just one longer non-coding intergenic sequence, of 196 base pairs.

As was already indicated by the 9.7kb published partial genome^25^, the gene arrangement we found in *P. rubra* is unique amongst published metazoan mitochondrial genomes. Similarly, *I. pulchra* shows no similarity to any other published metazoan mitochondrial genome (Fig. 5). The species analysed in this study share only the small 'block' of *nad3-atp6-nad4-cob* (*I. pulchra*) and *cob-nad4-nad3* (*P. rubra*). However, the order is reversed between the two, and the genes are distributed across both strands in *I. pulchra*, so it is unlikely that this represents a feature inherited from a common ancestor. To quantify the number of common gene arrangements between the species in this study and other mitochondrial genomes, protein-coding gene and ribosomal RNA gene order was analysed using CREx^28^ (compared to the acoel *S. roscoffensis*, the xenoturbellid *Xenoturbella bocki*, and the metazoan mitochondrial 'ground plan', represented by *L. polyphemus*). Conserved mitochondrial gene 'blocks' (that is, a series of genes, regardless of their order within the grouping) were very infrequent between the species. Of the genomes compared, the highest number of common gene blocks was found between *X. bocki* and *P. rubra*. This result was not significant, finding only 16 common intervals out of a possible 176, and confirming the visual observation that gene order between these species is highly variable.

**Figure 5:**
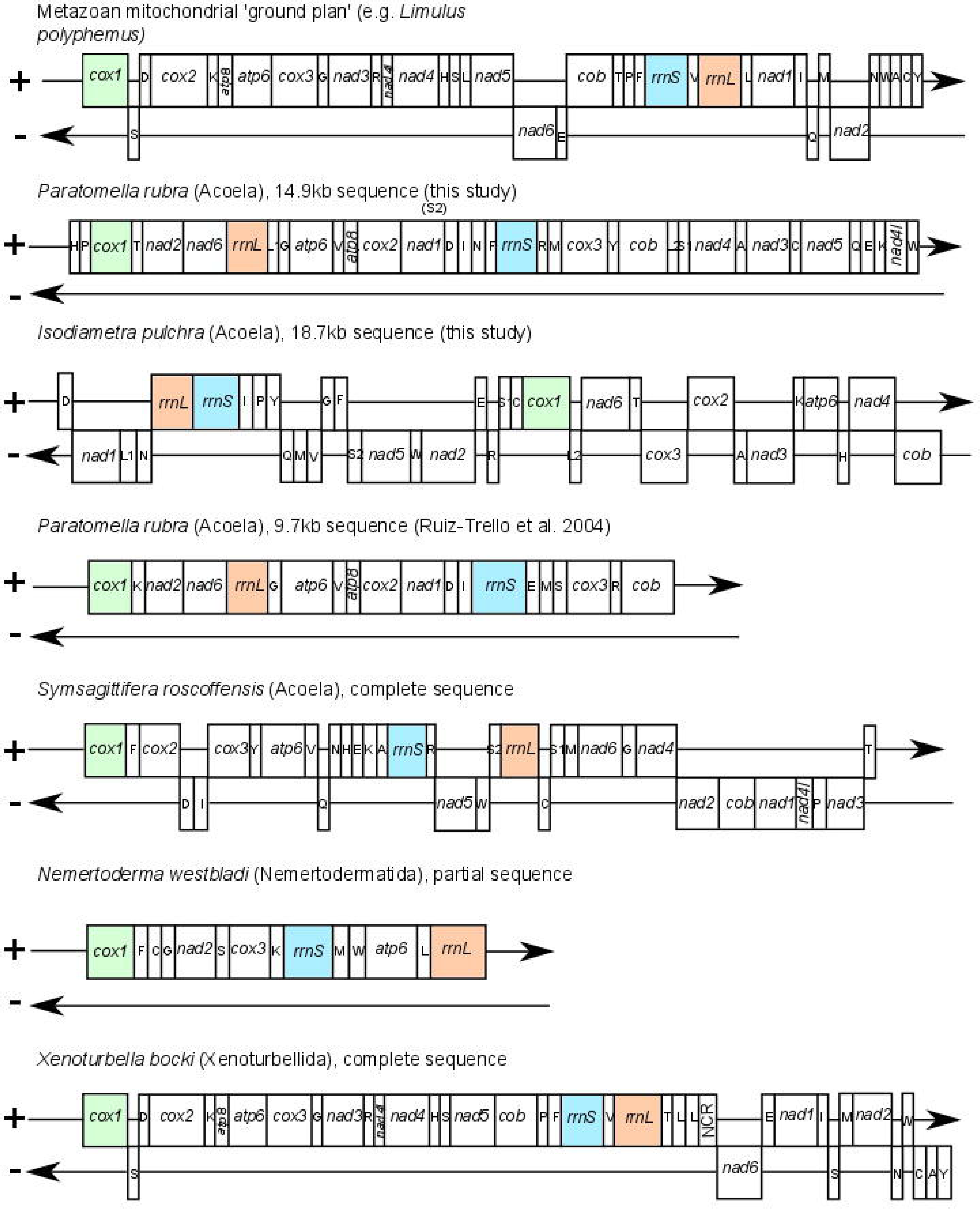
Comparisons of gene orders in the mitochondrial genome sequences resolved for *Paratomella rubra* and *Isodiametra pulchra*, compared to a published *P. rubra* fragment; the acoel *Symsagittifera roscoffensis*; the xenoturbellid *Xenoturbella bocki*; the nemertodermatid *Nemertoderma westbladi* and the metazoan mitochondrial 'ground plan' gene order, represented by *Limulus polyphemus.* Genes are not drawn to scale. Coloured genes chosen to show 'anchors' and divergence from the ground plan gene order in other species.

### Phylogenetic Analysis, and population differentiation

We used our new mitochondrial data from *P. rubra* and *I. pulchra* to investigate the internal phylogeny of the acoels and to test support for an Acoela-Xenoturbellida affinity. Bayesian phylogenetic inference was carried out using the protein-coding genes of *P. rubra* and *I. pulchra* in an amino acid alignment, using the additional species listed in Supplementary Table S3.

Our maximum likelihood approach grouped both *P. rubra* and *I. pulchra* inside Acoela, as expected (Fig. 6). However, the Acoela were clustered with Tunicata in our analysis and this acoel/tunicate grouping was found as an outgroup to all other Bilateria. The protostome/deuterostome split was correctly inferred and Xenoturbellida were found splitting off inside Deuterostomia. We detected the Nemertodermatida species *Nemertoderma westbladi* inside Mollusca and we therefore conclude that the limited data available on Genbank for this species is a contamination.

**Figure 6:**
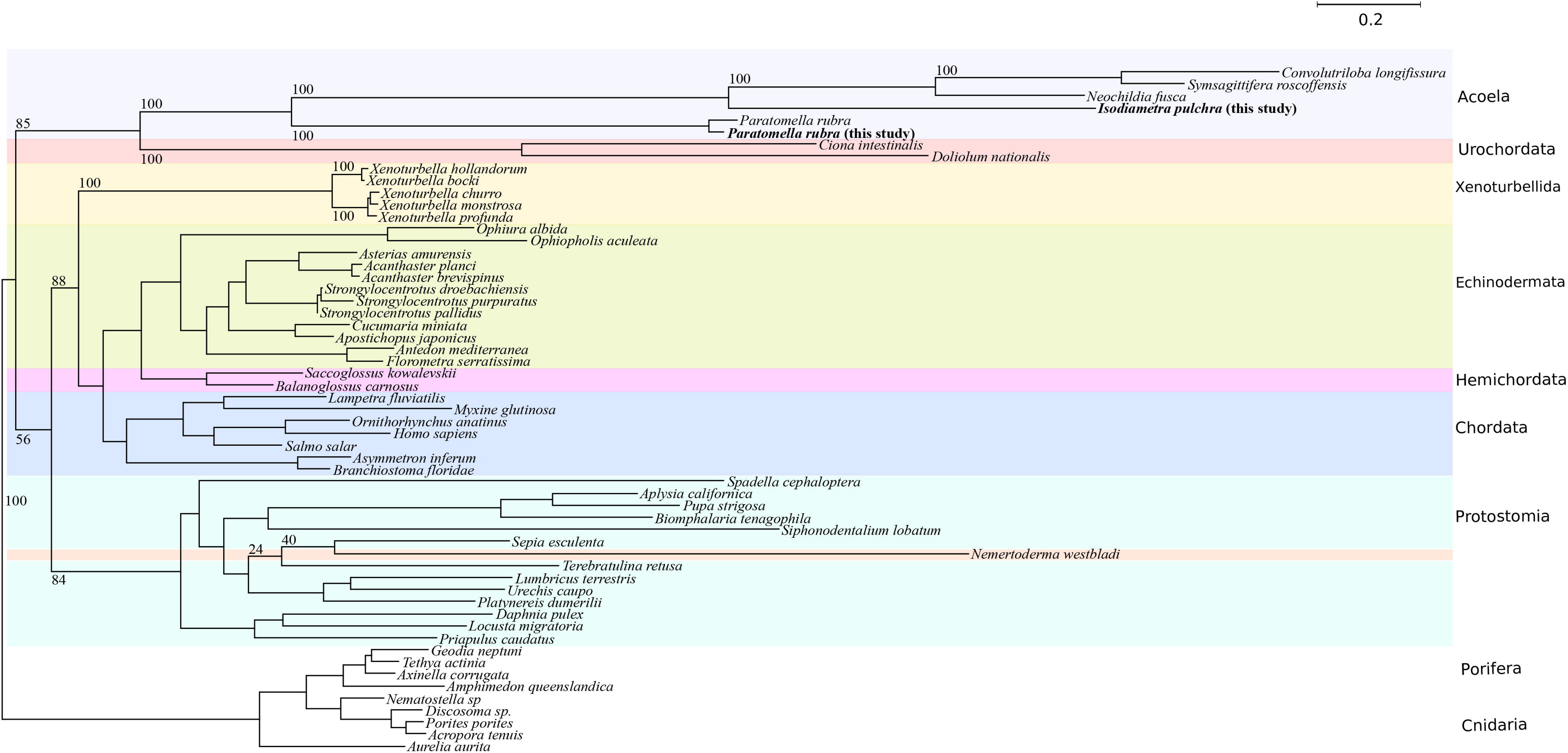
Maximum Likelihood (using RAxML) phylogenetic analysis of mitochondrial protein-coding genes from the Metazoa, including *P. rubra* and *I. pulchra*, with bootstrap support values at relevant nodes.

The Bayesian analysis of the untrimmed alignment did not converge. The concatenated and trimmed data set reached a maximum difference of 0.26 after 9921 trees were sampled in each of the 10 chains discarding the first 1000 trees (per chain) as burnin. While the grouping of Acoela and Tunicata is also found by this method with 96% posterior probability, the major clades within Bilateria (Chordata, Protostomia, Xenoturbellida, Hemichordata and Ambulacraria) remain unresolved in a polytomy with 60% posterior probability (Fig. 7).

**Figure 7:**
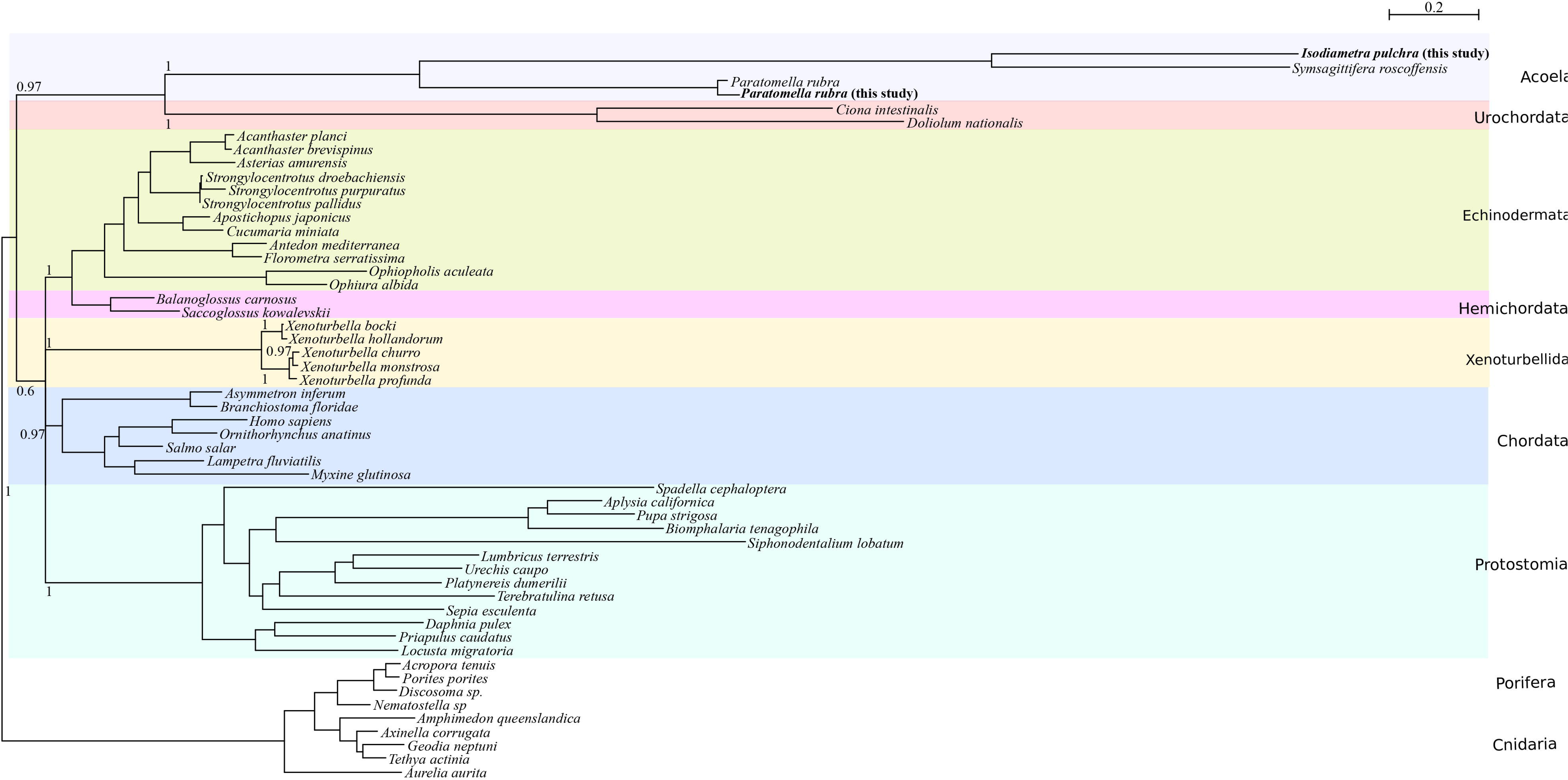
Bayesian (using PhyloBayes 50) phylogenetic analysis of mitochondrial protein-coding genes from the Metazoa, including *P. rubra* and *I. pulchra*, with posterior probabilities relevant nodes. Analysis carried out on trimmed alignment.

Having access to the published partial *P. rubra* mt sequence from a population sampled near Barcelona (Spain) and our own samples from Yorkshire, UK, we could estimate total sequence divergence and compare non-synonymous to synonymous substitutions in eight protein coding genes found on the Spanish fragment to the same genes from the Yorkshire mitochondrial genome. We found the 9.7kb sequences to be 82.62% similar at the nucleotide level. The number of substitutions to varied between, e.g. 23 in the shortest gene alignment (atp8; 177bp), to 161 in nad2 (972bp), and 116 in the 1401bp long cox1 alignment (Supplementary Table S2). Notably, non-synonymous substitutions appear to be frequent with, e.g. 13 in atp8, 104 in nad2, 25 in the cox1, see Supplementary Table S2.

## Discussion

### Genomic sequences

The 14.9kb sequence of the *P. rubra* mitochondrial genome determined in our analysis contains the full complement of 37 genes typical of metazoan mitochondrial DNA. Numerous lab-based and computational efforts to close the circular mitochondrial genome were unsuccessful, but the complete gene complement and length of our final sequence indicates that this fragment covers the majority of the *P. rubra* complete mitochondrial genome. The difficulty we encountered in attempting to close the circular mitochondrial sequence may be attributed to the AT-rich and repetitive sequence found at both ends of the fragment, which could have prevented PCR amplification. Similar regions have been shown as problematic in other studies of mitochondrial genomes^29^. As no long stretch of non-coding sequence was found for this species in our study, the missing sequence might represent its mitochondrial transcription control region. Nonetheless, the overall AT content of the *P. rubra* mitochondrial sequence (78.15%) is high even for mtDNA, and greater than the A+T content of the mitochondrial genome of the acoel *S. roscoffensis* (75.3%)^11^ and the published partial *P. rubra* genome (76.4%)^25^.

The validity of the duplicated sequence found in our analysis of the *I. pulchra* mitochondrial genome could not be confirmed by PCR or computational efforts to map short reads to resolve it. Duplications within mitochondrial genomes are not uncommon, and changes to mitochondrial gene order are widely thought to arise as a result of a sequence ‘duplication and deletion’ mechanism^14,30,31^. A number of mitochondrial genomes with duplicated sequences have been reported in species with a divergent mitochondrial gene order^31–33^. Given the highly unusual gene order of the *I. pulchra* mitochondrial genome, a genomic duplication such as this could provide evidence for a genomic 'duplication and deletion' rearrangement of genes. The rearrangement and separation of protein-coding genes in other mitochondrial genomes has been attributed to long, non-tandem, inverted repeats ^32^, and this could be true for *I. pulchra*. Furthermore, very long nematode mitochondrial genomes with variable duplicated regions have been found with a conserved region containing the majority of the protein-coding genes^34^: in *I.pulchra* the protein-coding genes and tRNAs, with the exception of *nad1* and *trnD* and *L1* are found in one long region, outside of the duplicated section. However, long non-coding duplications are frequently adjacent to tRNA*s* or other sequences capable of forming stem-and-loop structures ^35^. This is not true for the potential duplicate in *I. pulchra*. Most puzzling, both occurrences of the duplicate are identical, nucleotide-by-nucleotide, and unless the duplication occurred exceptionally recently, it is likely that spontaneous mutations would result in differences between the two copies of the sequence, especially given the elevated mutation rate of mitochondrial genomes. While it is true that the duplicated sequences appear at the start and end point of transcriptome assembly contigs, meaning it is possible that the duplication observed occurred only as a result of a sequencing and assembly error, their existence is nevertheless supported by PCR products which show an identical sequence being adjacent to both *rrnL* and to *cob*.

### Gene rearrangement: order and structure of the mitochondrial genome

The 14.9kb mitochondrial genome of *P. rubra* and the 18.7kb sequence from *I. pulchra* show no significant organisational similarity to any other published metazoan mitochondrial genome (Fig. 5). Comparison of the 14.9kb *P. rubra* sequence with the published 9.7kb *P. rubra* piece shows an identical protein-coding and ribosomal gene order, but with variation in tRNA order (Fig. 5). tRNAs are reported to show much more frequent gene translocation compared to larger genes, which could account for these discrepancies. This difference can be attributed to the different populations used in each study ^36^: the previous analysis used *P. rubra* collected from Barcelona, Spain^25^, whilst the animals used in this study came from Yorkshire, England.

Both species analysed in this study are unique in the orientations and orders of their genes: *P. rubra* has genes exclusively on one strand, whilst *I. pulchra* has an almost-equal distribution of genes across both the plus (18 genes) and minus (17 genes) strands. Furthermore, genes in *I. pulchra* are not clustered into groups of 'gene blocks' on the same strand, but are found frequently as one or two genes on each strand. The unique gene order found for these species seems to be typical for the acoels: analysis of the complete *S. roscoffensis* mitochondrial genome found no gene order similarity to any other species published to date, suggesting great variability in mitochondrial gene order amongst the acoels ^11^. Although mitochondrial gene order can be used as a tool for phylogenetic inference, the huge discrepancies in gene order amongst the acoels suggests that this cannot be an informative tool for this clade.

In addition to an unusual gene order, the mitochondrial genome of *P. rubra* shows frequent overlaps between protein-coding genes and tRNAs. tRNAs have been reported within protein-coding genes in other metazoan mitochondrial genomes ^37,38^, and given that no other location could be predicted for these sequences, this overlap could represent the simultaneous coding for both tRNAs and protein-coding genes. Overlap in coding sequence could be the result of a tendency to reduce genome size, accompanied by a reduction in non-coding sequence ^38^, and truncated tRNAs with incomplete secondary structure ^39^, both of which are also found for the *P. rubra* sequence. The opposite is true for the *I. pulchra* sequence. The sequence we could confidently verify makes the minimal length of the *I. pulchra* mitochondrial genome 18,725 base pairs, and likely to be longer in the complete closed circular genome. As has been found for other 'long' mitochondrial genomes, this is largely due to a large portion of the genome being non-coding ^40^. The lengths of protein-coding genes inferred for *I. pulchra* are similar to those of other acoel species (Table 3), and in addition, two protein-coding genes (*atp8* and *nad4l*) appear to have been lost from the genome, contributing to a reduced proportion of protein-coding gene sequence within the genome. The loss of *atp8* is not unusual, and has been reported in a number of unrelated taxa, as well as *S. roscoffensis* ^11^. The absence of *nad4l* is more unusual, and could be a result of transfer to the nucleus or its existence in a portion of the genome that we have been unable to sequence.

**Table 3:**
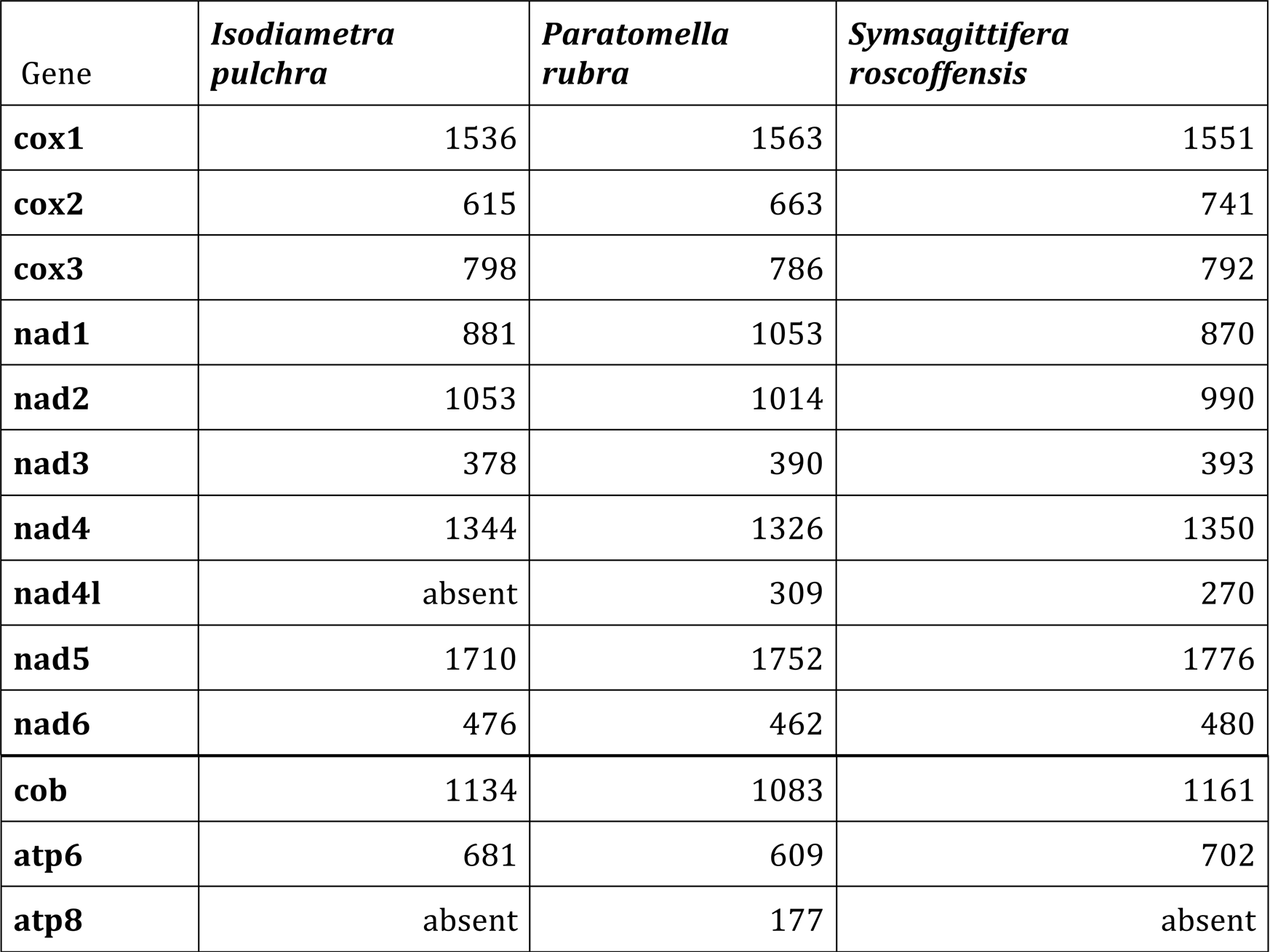
Length of protein-coding genes in acoel mitochondrial genomes. All gene lengths in base pairs.

| Gene | *Isodiametra pulchra* | *Paratomella rubra* | *Symsagittifera roscoffensis* |
| --- | --- | --- | --- |
| **cox1** | 1536 | 1563 | 1551 |
| **cox2** | 615 | 663 | 741 |
| **cox3** | 798 | 786 | 792 |
| **nad1** | 881 | 1053 | 870 |
| **nad2** | 1053 | 1014 | 990 |
| **nad3** | 378 | 390 | 393 |
| **nad4** | 1344 | 1326 | 1350 |
| **nad4l** | absent | 309 | 270 |
| **nad5** | 1710 | 1752 | 1776 |
| **nad6** | 476 | 462 | 480 |
| **cob** | 1134 | 1083 | 1161 |
| **atp6** | 681 | 609 | 702 |
| **atp8** | absent | 177 | absent |

### Acoela and Xenacoelomorpha phylogenetics

The internal phylogeny we resolve for Acoela is in line with that proposed by Jondelius et al. ^24^. *I. pulchra* groups with *S. roscoffensis*, *Neochildia fusca* and *Convolutriloba longifissura*, which are all members of the Convolutidae. *P. rubra*, forms a separate branch outside the Convolutidae representing the Paratomellidae. However, we do not retrieve Xenacoelomorpha, but find the acoels grouped with tunicates outside of the Bilateria as sister taxon to the non-bilaterian outgroups. The Xenoturbellida in our analysis are placed as a branch of the deuterostomes, as expected. We interpret the grouping of the acoels and tunicates as a classical example of long branch attraction (LBA) (Fig. 6), where the fast evolving mitochondrial sequences of both taxa are drawn towards each other and outgroup species. The accelerated substitution rates in mitochondrial DNA are also evidenced by the population divergence we find in *P. rubra*, and may well lead to LBA in phylogenies derived from mitochondrial protein-coding genes, owing to the clustering of rapidly evolving lineages. This is of particular relevance for acoel species, which already demonstrate a very rapid rate of nucleotide substitution compared to other metazoans, leaving them vulnerable to LBA and incorrect clustering in a position outside of protostomes/deuterostomes ^4,41^.

The mitochondrial genome sequences we analysed for the acoel species *P. rubra* and *I. pulchra* have very divergent gene orders compared to other metazoan species. Furthermore, both species have very different mitochondrial features: a large mount of genomic overlap in *P. rubra*, and a lot of non-coding sequence in *I. pulchra*. It is also possible that the mitochondrial genome of *I. pulchra* has a non-tandem inverted duplication - which could provide a mechanism for gene order variation - but this could not be confirmed by lab or computational based methods. Although limited to three species, the uniqueness of acoel mitochondrial genomes analysed so far ^11,25^, means that gene order and other mitochondrial genome features may not be phylogenetically informative for this order, although further mitochondrial genomes from other members of the Acoela would no doubt aid in this comparative analysis. Our data clearly emphasise the still problematic placing of Xenacoelomorpha, with the *Xenoturbella* clade firmly placed inside deuterostomes, but LBA drawing the acoels (and the tunicates) towards the outgroup. In summary, more data from genomes of early branching taxa is thus needed to resolve phylogenetic and biological questions.

## Methods

### Specimen Collection, DNA extraction and PCR

Live *Paratomella rubra* specimens were isolated from sand samples collected from Filey, North Yorkshire and were immediately frozen and stored at −70°C following identification. Specimens of *Isodiametra pulchra* were cultured in petri dishes with nutrient-enriched f/2 sea water and fed ad libitum on *Nitzschia curvilineata* diatoms. DNA was extracted from live specimens of *I. pulchra* and frozen specimens of *P. rubra* using the QIAamp DNA Micro Kit (Qiagen: Product No. 56304) with the manufacturer recommended protocol.

All PCRs were done using the GeneAmp PCR System 2700 (Applied Bioscience). PCRs were carried out using the Expand Long-Range PCR Kit (Roche Applied Sciences: Product No. 11681834001), following manufacturer recommendations for 50μl reaction set-up. General cycling protocol was: 92°C for 2 min; 15 cycles of: 92°C for 10 sec, 57°C for 15 sec, 68°C at initial elongation time (approximated as 1 min per 1000 base pairs to be amplified); 2 cycles each of: 92°C for 10 sec, 57°C for 15 sec, 68°C at 40 sec longer than initial elongation time, repeated at increasing 40 sec intervals for a further 14 cycles; a final elongation stage at 68°C for 7 min and a 4°C ‘hold’ stage. Where PCRs were not successful using this protocol, they were repeated using the Q5 High-Fidelity PCR Kit (New England Biolabs: Product No. E0555L), following manufacturer recommendations for a 25μl reaction. Cycling protocol was: 92°C for 2 min; 40 cycles of: 92°C for 10 sec, 58°C for 15 sec, 68°C at initial elongation time (approximated as 1 min per 1000 base pairs to be amplified); a final elongation stage at 68°C for 7 min. Amplified products were visualised on ethidium-bromide stained TAE 0.8% gels. Bands of expected size were purified using the High Pure PCR Product Purification Kit (Roche Applied Sciences: Product No. 11732668001) and sent for sequencing by Source BioScience Life Sciences. Only amplifications that resulted in one clear band on the TAE 0.8% agarose gel were sequenced.

Three fragments of sequence from the mitochondrial genome of *P. rubra*, of size ~5.8kb, ~4kb, and ~1.2kb, were generated from our gDNA assembly. Fragments were verified using a translated nucleotide query blast with invertebrate codon usage (blastx NCBI), and their orientation determined by gene annotation in comparison to the published 9.7kb section of the *P. rubra* mitochondrial genome ^25^. Primers were designed in conserved gene regions to:

1) Amplify across the ‘N-stretches’ present in the 5.8kb and 4kb fragments (8 and 9 N-stretches respectively, all of arbitrary length 50 base pairs)

2) Cover the whole 1.2kb fragment, with the aim of resolving the two frameshift mutations within the assembled sequence

3) Close the circular mitochondrial genome, by joining the 5.8kb fragment to the 1.2kb fragment; the 1.2kb fragment to the 4kb fragment; and the 4kb fragment to the 5.8kb fragment. (See Supplementary Figure S1). Amplification of the fragments joining the 1.2kb fragment to the 4kb fragment and to close the mitochondrial genome using standard PCR cycling were unsuccessful. These were repeated using a touchdown protocol with Expand Long-Range polymerase. Annealing temperature was set at 65°C with decreasing 2°C intervals every 2 cycles down to 49°C. Initial elongation time was calculated as before, increasing 30 sec every two cycles of the touchdown, with a final 6 cycles at 49°C. This successfully amplified the region joining the 1.2kb fragment to the 4kb fragment, but we could not close the circular genome. Design of three new forward and reverse primers, tried in all combinations and using variable PCR parameters were unsuccessful in closing the mitochondrial genome. Additional RNA-seq and DNA genomic sequencing data corroborated the stretches of sequence at either end of the mitochondrial genome but did not aid in closing the circle.

Three mitochondrial contigs of size ~13kb, ~3.5kb and ~1.3kb were identified from *I. pulchra* Trinity transcriptome assembly from total RNA sequencing. A further contig of ~19kb was also identified, covering the entire ~1.3 and 13kb regions, and ~2.4kb of the 3.5kb sequence. Fragments were verified using blastx, NCBI, as outlined for *P. rubra*, and approximations for the location of protein-coding genes and tRNAs determined using the MITOS mitochondrial genome annotation server^42^ (http://mitos.bioinf.uni-leipzig.de/help.py). Primers were designed to span the 13kb contig in two ~5kb sections, and to join the 13kb contig to the 3.5kb contig in both directions, to close the mitochondrial genome and check the validity of the duplicated region. (Fig. 2). RNA-Seq data for *I. pulchra* were mapped to the long transcriptome assembly contigs and PCR sequencing results using NextGenMap ^43^, and visualised using Tablet ^44^.

### Genome Annotation

For *P. rubra*, all sequenced fragments were aligned against the initial scaffold of the 9.7kb published sequence^25^; the 5.8kb, 4kb and 1.2kb genome assembly sequences; and an additional long genome assembly fragment of length 14,954 (see Supplementary Figure S1). All contigs and PCR sequencing results were similarly aligned for *I. pulchra*, but without a reference sequence (see Fig. 2). Alignments were visualised using Mesquite (http://mesquiteproject.org). with invertebrate mitochondrial translated amino acid state colour coding. Where ambiguity remained between PCR sequencing results and genome or transcriptome assembly fragments, the genome or transcriptome assembly nucleotide sequence was used to establish a final 'consensus' sequence for each mitochondrial genome. This was of particular relevance for repetitive AT regions – for example, within *P.rubra nad1*, for which PCR sequencing results were inconclusive. In the case of *I. pulchra*, where the validity of the duplicated sections could not be confidently determined, separate consensus sequences were made: one covering the region from the 'start' of the long 19kb contig to the first occurrence of the duplicated sequence; one covering the region from the end of the first duplicate to the start of the second duplicate, and one 'long' closed-circle sequence, containing both occurrences of the duplicated sequence.

The region for each protein-coding mitochondrial gene (*nad1-6, nad4l, cox1-3, cob, atp6 and atp8*) in the new consensus sequence was compared against published mitochondrial genomes using a translated nucleotide query (blastx, NCBI) with NCBI translation table number 5 ‘Invertebrate mitochondrial’. Published genes from the mitochondrial genomes of the acoels *Symsagittifera roscoffensis* and *P.rubra* were downloaded from the NCBI Nucleotide database and aligned to the new consensus gene sequences of both *P. rubra* and *I. pulchra* to verify the location of protein-coding and ribosomal RNA-encoding genes. The 5’ end of protein-coding genes were inferred to start from the first in-frame start codon (ATN, GTG, TTG, or GTT), even if this appeared to overlap with the preceding gene. Similarly, the terminal of protein-coding genes was inferred to be the first in-frame stop codon (TAA, TAG, or TGA). If no stop codon was present, a truncated stop-codon (T-- or TA-) prior to the beginning of the next gene was assumed to be the termination codon, completed by post-transcription polyadenlylation. tRNA sequences and putative secondary structures were identified using the Mitfi program within MITOS.

### Sequence Alignment, Phylogenetic Analysis, and evolutionary rates

Phylogenetic analysis was performed using a concatenated amino acid alignment of all thirteen protein-coding genes for *P. rubra*, and all eleven protein-coding genes present in *I. pulchra*. Nucleotide sequences were aligned using TranslatorX (http://www.translatorx.co.uk/) independently for all genes with invertebrate mitochondrial genetic code, using ClustalOmega ^45^ for amino acid alignment. Protein alignments were reduced to the most informative residues using trimAl v.1.4.rev15^46^ with standard settings. Predicted protein-coding amino acid sequences from both taxa were imported into an aligned matrix comprising 54 other species, taken from a wide range of published metazoan mitochondrial genomes representing deuterostomes, protostomes, cnidarians, and two species of poriferans as an outgroup (Supplementary Table S3). Regions showing ambiguity in alignment were excluded, so that only blocks of well-aligned sequence were included for analysis.

We initially re-constructed neighbour nets in SplitsTrees v.4^47^ to screen our dataset for potentially non-treelike patterns, which could impede our phylogenetic analysis. Subsequently, we used RAxML v. 8.2.9^48^ to infer maximum likelihood phylogenies from the original and the reduced alignments under the MTZOA model^49^. Bootstrapping was conducted employing the "autoMRE” option in RAxML and the trees visualised with figtree v.1.4 (http://tree.bio.ed.ac.uk/software/figtree). We carried out Bayesian inference on the trimmed alignment with PhyloBayes v.4.1^50^ under the MTZOA model. We ran 10 chains in parallel and stopped the tree search at 9921 per chain, with a maximum difference of 0.26, when discarding 1000 trees as burnin and sampling every 10th tree per chain.

We used the Geneious software (v.8) to calculate sequence differences for a 9.7kb section of the *P. rubra* mitochondrial genome originating from worms sampled in Filey, Yorkshire (UK) and Barcelona (Spain)^25^. For eight protein-coding genes found on this section we used ParaAT (v2.0)^51^ to calculate translation alignments and the KaKs calculator (v1.2)^52^ to access substitution rates.

## Acknowledgements and Funding

This work was supported by the European Research Council (ERC-2012-AdG 322790), the Leverhulme Trust (grant F/07 134/DA) and the Biotechnology and Biological Sciences Research Council (grant BBS/H006966/1). MJT was supported by a Royal Society Wolfson Research Merit Award.

We declare that no competing interests exist.

We would like to thank Richard Copley (The Wellcome Trust Centre for Human Genetics, University of Oxford) and the Oxford Wellcome Trust for their assistance with *P. rubra* sequencing, We are indebted to Anne Zakrzewski for her assistance with animal collection trips, associated animal identification, and valuable comments on the manuscript.

### Authors contributions

Conceived and designed the experiments: HER FL BE. Performed the experiments: HER, BE, FL, PHS. Analysed the data: HER PHS MJT. Wrote the paper: HER, MJT, PHS.

